# A multivariate to multivariate approach for voxel-wise genome-wide association analysis

**DOI:** 10.1101/2021.11.02.467021

**Authors:** Qiong Wu, Yuan Zhang, Xiaoqi Huang, Tianzhou Ma, L. Elliot Hong, Peter Kochunov, Shuo Chen

## Abstract

The joint analysis of imaging-genetics data facilitates the systematic investigation of genetic effects on brain structures and functions with spatial specificity. We focus on voxel-wise genome-wide association analysis, which may involve trillions of single nucleotide polymorphism (SNP)-voxel pairs. We attempt to identify underlying organized association patterns of SNP-voxel pairs and understand the polygenic and pleiotropic networks on brain imaging traits. We propose a *bi-clique* graph structure (i.e., a set of SNPs highly correlated with a cluster of voxels) for the systematic association pattern. Next, we develop computational strategies to detect latent SNP-voxel *bi-cliques* and inference model for statistical testing. We further provide theoretical results to guarantee the accuracy of our computational algorithms and statistical inference. We validate our method by extensive simulation studies, and then apply it to the whole genome genetic and voxel-level white matter integrity data collected from 1052 participants of the human connectome project (HCP). The results demonstrate multiple genetic loci influencing white matter integrity measures on splenium and genu of the corpus callosum.

## 1 Introduction

Imaging-genetics has garnered increased interest in the field of neuropsychiatric research as it provides a viable pathway to understand brain diseases by integrating genetic, brain imaging, and environmental factors. The joint analysis of imging-genetics data reveals the genetic effects on spatially specific brain functions and structures (Ge et al., 2013; Liu and Calhoun, 2014; Nathoo et al., 2019; Smith et al., 2021; Zhao et al., 2019, 2021; Zhu et al., 2014). Identifying genetic effects on objectively measured high-resolution imaging traits can enhance understanding the complex genetic and neurological mechanisms of neuropsychiatric disorders.

In imaging-genetics studies, both brain imaging data and genome sequence are measured for each participant. The genetic measurements can characterize genetic variations using single nucleotide polymorphism (SNP) and copy number variants (CNVs). The non-invasive brain imaging techniques assess the brain structures by magnetic resonance imaging (MRI), diffusion tensor imaging (DTI), and brain functions by functional magnetic resonance imaging (fMRI). The recent development of neuroimaging technology provides high-resolution imaging data with improved spatial specificity and thus can better assess the genetic effects on brain structures and functions.

The statistical analysis of imaging-genetics data is computationally intensive and methodologically challenging. These challenges mainly rise from the combination of two sets of high-dimensional features: multivariate imaging traits with multivariate genetic variants. Moreover, both imaging traits and genetic variants exhibit complex and organized dependence structure reflecting the underlying neurophysiological mechanisms and linkage disequilibrium patterns (Nathoo et al., 2019). For example, a typical imaging-genetics study collects up to 10^7^ SNPs and 10^5^ voxels, jointly contributing trillions (10^12^) of SNP-voxel pairs (Huang et al., 2015, 2017). The direct application of classic voxel-wise genome-wide association analysis (vGWAS) may require an enormous sample size (e.g., multiple millions of participants) to control the false positive error rate while maintaining adequate statistical power (Ge et al., 2012, 2015; Hibar et al., 2011; Stein et al., 2010).

Furthermore, advanced methods have been developed to leverage group sparsity by techniques including regularization, low rank techniques and projection of high-dimensional features (Chi et al., 2013; Greenlaw et al., 2017; Hardoon et al., 2009; Kong et al., 2020; Le Floch et al., 2012; Liu et al., 2009; Wang et al., 2012; Vounou et al., 2010, 2012; Zhu et al., 2014). However, while these methods could gain statistical power by jointly modelling genetic variants and imaging traits through a multivariate regression model, the high dimensionality of imaging-genetics data remains challenging due to computational burdens and/or overfittings. The results from summarized measures as a few latent variables or a coarser scale are less interpretable or lacking the spatial specificity (Liu and Calhoun, 2014).

In this study, we propose a new multivariate to multivariate method to systematically investigate the SNP-voxel association patterns with four aims: identify voxel clusters as genetically correlated imaging traits, detect functionally related SNP sets, understand the SNP-voxel association patterns as polygenic and pleiotropic relationships, and test the association patterns while controlling multiplicity. Specifically, we consider genetic variants and imaging voxels as two disjoint sets of nodes, correspondingly, and associations between all SNP-voxel pairs as edges in a bipartite graph. We model the polygenic and pleiotropic SNP-voxel association structure as an imaging-genetics *dense* bi-clique (IGDB). IGDB is a node-induced subgraph consisting of a subset of SNPs and a subset of voxels, where the possibility of a SNP associated with a voxel is much elevated than the rest of graph. Within an IGDB, each voxel can be considered as a polygenic imaging trait, and a SNP as a pleiotropic genetic variant. The existence of the polygenic and pleiotropic SNP-voxel association structure can be evaluated against a random bipartite graph. We then develop computationally efficient algorithms to extract the IGDB structure from the bipartite graph mixture model and thus provide sound estimates of parameters in the mixture model. Our inference on IGDB is constructed via likelihood based statistic on the bipartite graph mixture model, and thus can improve statistical power with controlled family-wise error rate.

## 2 Motivating Data Example

The Human Connectome Project (HCP) sponsored by National Institutes of Health (NIH) aims to construct the underlying neuro pathways with healthy human brain functions. It is an important public resource for structural and functional brain connectivity data, accompanied by demographic, behavioral, genetic and other data. In this study, we focus on the brain imaging and genetics data in the HCP surveyed from 1052 participants (F/M 483/569; age 28.1±3.7), for whom the scans and data were released in June 2014 (https://humanconnectome.org) that passed the HCP and ENIGMA quality control and assurance standards (Marcus et al., 2013). The participants in the HCP study were recruited from a large population-based study named “the Missouri Family and Twin Registry” (Van Essen et al., 2013).

The fractional anisotropy (FA) measure, derived from diffusion tensor imaging (DTI), is a widely-used brain structural connectivity metric for studying the white matter microstructure. Previous studies have investigated the heritability quantitatively through variance components method of pedigrees (Jahanshad et al., 2013; Kochunov et al., 2014). They find that 70% to 80% of the total phenotypic variance of tract-wise FA measures can be explained by additive genetic factors (Kochunov et al., 2015). The significantly and reliably hertiable FA measurements are qualified as a set of endophenotypes which suggests to further specify genetic variants associated with these traits. Hence, the genetic analysis is desirable to detect the genetic effect from specific loci on imaging traits with statistical inference. Moreover, it is reported that FA measurements at multiple brain locations can be affected by a common set of genetic variates (Zhao et al., 2021). FA is a complex trait determined by multiple alleles. It stimulates the identification of functionally-related genetic variants. This investigation naturally invokes the search for polygenity and pleiotropy networks as the focus of this study. Voxel-level association analysis between imaging traits and genetic variants can provide the maximal spatial resolution. Nevertheless, the implementation is challenging because it requires a multivariate to multivariate association analysis to extract SNP-voxel subnetworks with polygenic and pleiotropic structures and further to provide sound statistical inference. To close this gap, we develop an IGDB-based framework to perform voxel-vise GWAS and systematically identify polygenic and pleiotropic structures.

## 3 Methods

### 3.1 Background and notations

We consider an imaging-genetics data set collected from *L* independent subjects. We let *V* be the set of brain imaging voxels with |*V*| = *n* and *U* be the set of genetic variants (i.e., SNPs) with |*U* | = *m*. For each participant *l* ∈ {1, …, *L*}, define ***x***_*l*_ = (*x*_1,*l*_, …, *x*_*m,l*_)^*T*^ to be the genetic variants for the participant *l* and ***y***_*l*_ = (*y*_1,*l*_, …, *y*_*n,l*_)^*T*^ to be the vector of multivariate imaging traits. Let *z*_*l*_ denote a *p*-dimensional vector of individual-level profiling covariates We model the associations between multivariate imaging traits and multivariate genetic variants using a generalized linear regression model:

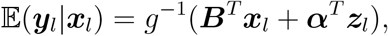

where *g*(·) is a known link function with inverse *g*^−1^(·), and the coefficient ***B*** = {*β*_*uv*_}_*u*∈*U,v*∈*V*_ ∈ ℝ^*m×n*^ is called the SNP-voxel association matrix. The goal of our statistical inference is to accurately identify the subset of significant associations {(*u, v*) : *β*_*uv*_ ≠ 0} based on multi-variate to multivariate hypothesis testing (Benjamini and Hochberg, 2000; Efron, 2012):

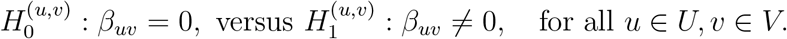

Conventional statistical inference methods (e.g., multiple testing correction or regression shrinkage) work by regularizing vectorized ***B***. However, this strategy may only capture individual association pairs *β*_*uv*_ without recognizing systematic patterns (e.g., the pleiotropic and polygenic structure). A prominent example is that a cluster of SNPs may jointly influence the observations on a cluster of neighboring voxels. To address this challenge, we propose a new multivariate to multivariate inference framework that extracts the joint structure in ***B***, which we call *imaging-genetics dense bi-clique (IGDB)*. Next, we introduce the IGDB structure, based on which, we then formally propose a novel estimation and inference procedure on this structure.

### 3.2 IGDB in a multivariate to multivariate graph structure

We characterize the vGWAS association as a bipartite graph *G* = (*U, V, E*), where *U* and *V* are distinct node sets representing SNPs and voxels, respectively. The set of binary edges *E* describes the locations of significant SNP-voxel associations: *e*_*uv*_ ∈ *E* if and only if *β*_*uv*_ ≠ 0 in the association matrix ***B*** = {*β*_*uv*_}_*u*∈*U,v*∈*V*_. In contrast to conventional approaches that treat edges *e*_*uv*_ individually, our proposal provides a succicint description of pleiotropic (one SNP to multiple image voxels) and polygenic (multiple SNPs to one voxel) relationships. To this end, we now formally propose IGDB as a subgraph structure of *G*. Denote an arbitrary subgraph of *G* by *G*[*S, T*] = (*S, T, E*[*S, T*]), where *S* ⊂ *U* , *T* ⊂ *V* and *E*[*S, T*] = {*e*_*uv*_ ∈ *E*|*i* ∈ *S, j* ∈ *T*}. Our proposed IGDB will be defined based on some particular subgraph *G*[*S*_0_, *T*_0_] such that most *β*_*uv*_’s are nonzero for *e*_*uv*_ ∈ *G*[*S*_0_, *T*_0_], while most *β*_*u′v′*_’s elsewhere are zero. Our core intuition can be quantified into the following formulation:

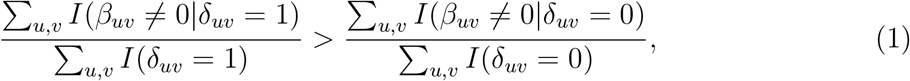

where *δ*_*uv*_ is a binary variable indicating the IGDB-based network structure, i.e.,

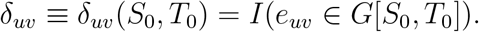

This reflects that imaging features (*T*_0_) are polygenic traits and the genetic variants (*S*_0_) are pleiotropic alleles. The genetically correlated imaging features and functionally related SNPs jointly compose a functional biclique *G*[*S*_0_, *T*_0_]. In neuroimaging studies, findings are often reported for spatially contiguous brain areas (i.e., connected voxels) because of the biological interpretability and inference advantages (Woo et al., 2014). This is reflected in our proposed IGDB structure by further formulating *S*_0_ and *T*_0_ as disjoint vertex neighborhoods, as follows:

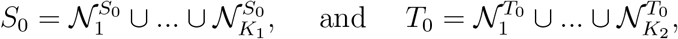

where each 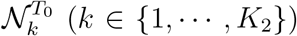 is a spatially contiguous voxel cluster, and accordingly 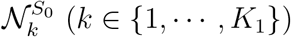 is a set of functionally related SNPs associated with one or multiple spatially-contiguous voxel clusters (e.g., 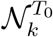). In the next subsection, we articulate that the IGDB enjoys several statistical advantages supported by graph and combinatorics theory.

### 3.3 Graph properties of IGDB

Without loss of generality, we consider the following two cases regarding the underlying network structure of *G*:

**Case 0** : *G* is observed from a random bipartite graph *G*(*m, n, μ*_0_),

**Case 1** : There exists at least one non-trivial IGDB *G*[*S*_0_, *T*_0_] such that *G* is observed from

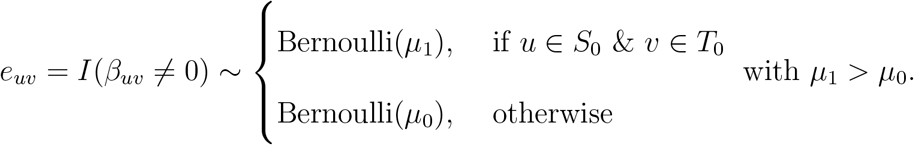

In Case 0 (i.e., no polygenic and pleiotropic patterns), we can directly implement the conventional multiple testing corrections and regression shrinkage methods to determine individual associations between genetic variants and imaging traits. If Case 1 presents, our primary goal becomes to extract and test the underlying IGDB subgraphs as polygenic and pleiotropic subnetworks.

In practice, the estimated IGDB from a sample can be used to distinguish Case 0 versus Case 1 because the observed network behave differently under two cases on the size of the maximal “dense” subgraph. For convenience, we call a subgraph *G*[*S, T*] a *γ-quasi biclique*, if it contains at least *γ* · |*S*| · |*T* | edges. Then, asymptotically, if |*S*_0_|, |*T*_0_| → ∞ as *m, n* → ∞, with high probability, the true IGDB subgraph *G*[*S*_0_, *T*_0_] would be a *γ*-quasi biclique for any fixed *γ* ∈ (*μ*_0_, *μ*_1_). In contrast, under Case 0, there would rarely exist a *γ*-quasi biclique of decent size with high density as the following lemma.

#### Lemma 1

*Suppose G is observed from a random bipartite graph G(m, n, μ_0_) as Case 0. G[S, T] is any subgraph with edge density* 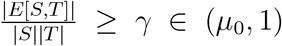 *(i.e., γ-quasi biclique). Let m*_0_, *n*_0_ = Ω(max{*m*^*ϵ*^, *n*^*ϵ*^}) *for some* 0 < *ϵ* < 1. *Then for sufficiently large m, n with c*(*γ, μ*_0_)*m*_0_ ≥ 8 log *n and c*(*γ, μ*_0_)*n*_0_ ≥ 8 log *m, we have*

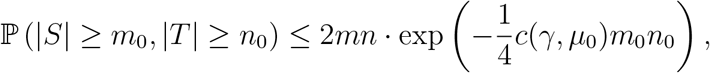

*where* 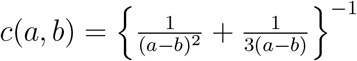.

## 4 Estimation and Inference

Let ***W***_*m*×*n*_ denote the inference result matrix (e.g., test statistics *w*_*uv*_ = *t*_*uv*_ or − log(*p*_*uv*_)) for the regression coefficients 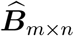. Then, our goal becomes to extract and test the IGDB structure from a weighted bipartite graph *G* = (*U, V*, ***W***). Similar to Efron (2012), as a natural consequence of our model set up in Section 3.2, edge weights in ***W*** follow a mixture marginal distribution:

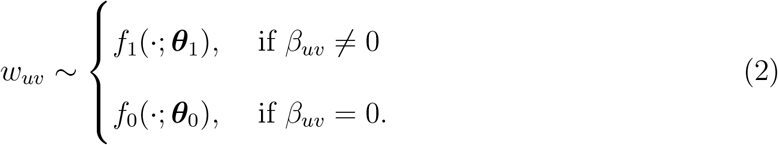

where *w*_*uv*_|*δ*_*uv*_ = 1 ∼ *μ*_1_*f*_1_ + (1 − *μ*_1_)*f*_0_, while *w*_*uv*_|*δ*_*uv*_ = 0 ∼ *μ*_0_*f*_1_ + (1 − *μ*_0_)*f*_0_. Empirically, we have the central tendency of 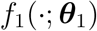 being greater than 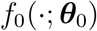, in the sense that 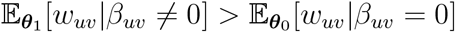.

## 4.1 IGDB estimation

Motivated by the nature of IGDB as a subgraph of elevated mean edge weights, we estimate it by looking for the maximal subgraph of *G* with a density constraint. Inspired by Lemma 1, we estimate the IGDB *G*[*S*_0_, *T*_0_] based on the edge weight matrix ***W*** by optimizing:

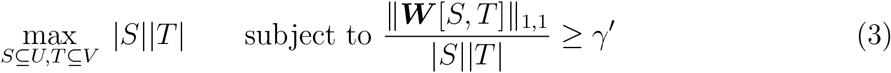

or the Lagrangian form after taking logarithm on both terms:

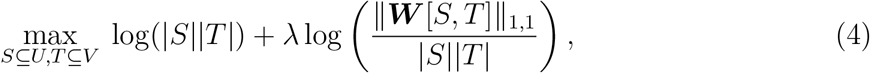

where || · ||_1,1_ refers to the entry-wise *ℓ*_1_ norm such that ||***W*** [*S, T*]||_1,1_ = Σ_*u*∈*S,v*∈*T*_ |*w_uv_*|, *γ*′ is the density constraint and the tuning parameter *λ* ∈ (1, ∞).

The direct optimization of the objective function (4) is challenging because it is a nondeterministic polynomial (NP) problem (Charikar, 2000; Khuller and Saha, 2009). We propose a computationally efficient greedy algorithm to approximately carry out the optimization of (4). We describe the greedy algorithm as Algorithm 1 in the following. In designing it, we extended the greedy algorithms for dense subgraph discovery (Khuller and Saha, 2009) in an adjacency matrix to a large bipartite matrix to extract dense bi-cliques. The computational complexity of Algorithm 1 is *O*(*C*_1_*mn*), where *C*_1_ is determined by the grid search of *h* (i.e., |*S*|/|*T*|) in the following Algorithm 1.

**Algorithm 1.**
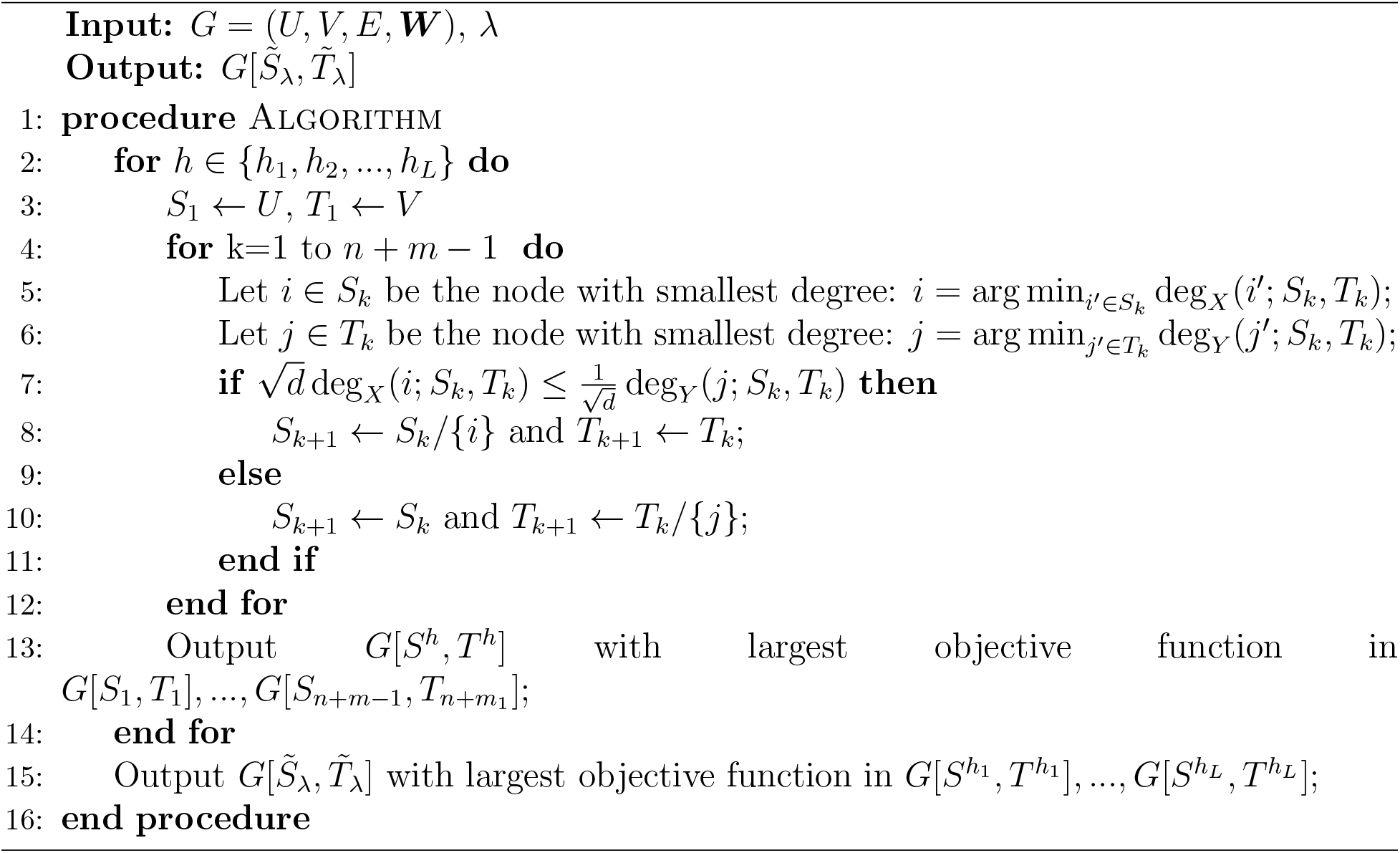
Direct optimization of objective function (4)

Now we establish approximation accuracy results of Algorithm 1 and its estimation of IGDB. Let 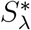 and 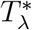 be the true optimal solution to (4):

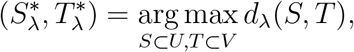

and 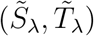 is from Algorithm 1 with

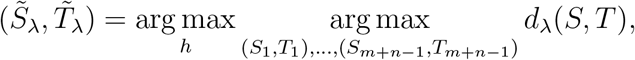

where 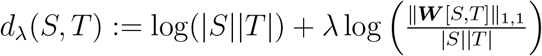.

The greedy algorithm with average-degree based density (or equivalently *λ* = 2) is said to have a 2-approximation guarantee for the true optimal (Charikar, 2000), namely, 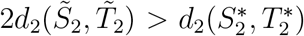. In this article, we present the approximation bounds for the proposed objective function (4) in terms of a parameter *λ* as the following Theorem 1.

### Theorem 1.

*For a given bipartite graph G* = (*U, V, E*), *with* 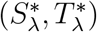) *and* 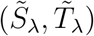 *defined in Section 3.1.1, the greedy algorithm 1 has a ρ*(*λ, m, n*)-*approximation, i.e.*, 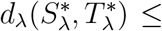 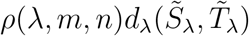 *with*

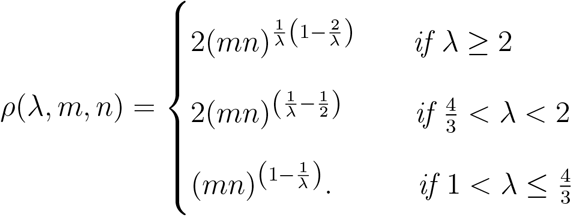

In Theorem 2, we state that the optimization of the proposed objective function (4) asymptotically leads to almost full recovery of the IGDB-based network structure.

### Theorem 2.

*Assume the graph G* = (*U, V, E*) *with an IGDB G*[*S*_0_, *T*_0_] = (*S*_0_, *T*_0_, *E*[*S*_0_, *T*_0_]) *is generated from mixture of Bernoulli distributions: e*_*uv*_ ~ *δ*_*uv*_*Bernoulli*(*π*_1_)+(1−δ_*uv*_)*Bernoulli*(*π*_0_), *δ*_*uv*_ = *I*(*e*_*uv*_ ∈ *G*[*S*_0_, *T*_0_]) and π_1_ > *π*_0_. *For simplicity, we let m* = *Θ*(*n*). *Assume* |*S*_0_| = *O*(|*m*|^1/2+*ϵ*^) and |*T*_0_| = *O*(|*n*|^1/2+*ϵ*^) *as n* → ∞ *for some ϵ* > 0. *Denote*

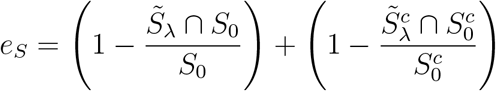

*and*

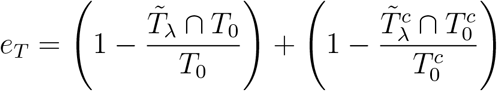

*to be the error rates of node memberships based on* 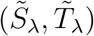 *from* Algorithm 1. *Then, there exists some λ such that we will get almost full recovery in* Algorithm 1, *i.e. for any fixed a* ∈ (0, 1), *as n* → ∞, *we have*

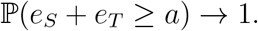

In practice, the tuning parameter *λ* can be objectively selected by a likelihood method (see the web Appendix A for details). Based on each dense subgraph *G*[*S, T*], we further identify spatially-contiguous voxel clusters (i.e., 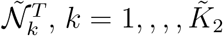, and a corresponding set of SNPs (i.e., 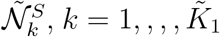) that are functionally associated with voxel clusters (see Web Appendix A). Last, multiple IGDBs can be extracted by performing algorithms repeatedly with the detected IGDBs masked (Cheng and Church, 2000).

### 4.2 Statistical inference of the IGDB

Recall that the purpose of this study is to perform statistical inference on the pleiotropic and polygenic association pattern or the IGDB. We investigate the significance of the presence of an IGDB against a random bipartite graph (Case 1 vs. Case 0) as illustrated in Section 3.3. Let *r* be a sound cutoff that dichotomize the weighted graph *G* into a binary graph *G*^*r*^ = (*U, V*, ***A***) using *a*_*uv*_ = *I*(|*w*_*uv*_| > *r*). Then, under IGDB structure indexed by node sets (*S*_0_, *T*_0_), the edges in *G*^*r*^ follow a mixture of two Bernoulli distributions:

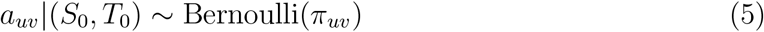

where *π*_*uv*_ = *δ*_*uv*_π_1_ + (1 − *δ*_*uv*_)*π*_0_ with 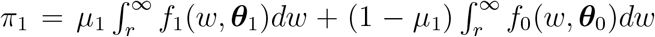, 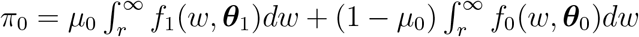 and *π*_1_ > *π*_0_. Then, a hypothesis testing to distinguish Case 0 and Case 1 can be proposed:

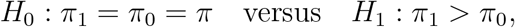

based on our mixture distribution model (5).

We propose a likelihood-based statistic for the IGDB test. For a binarized graph *G*^*r*^, let

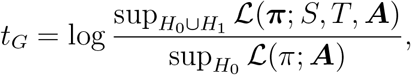

with likelihood given by Bernoulli distributions in (5). Then, the asymptotic power is ensured using the likelihood-based statistic through the following Theorem 3.

#### Theorem 3.

(Under IGDB alternative hypothesis H_1_). *Assume m* = Θ(*n*) *and the underlying IGDB G*[*S*_0_, *T*_0_] *with generating probabilities π*_1_ > *π*_0_ *satisfies* |*S*_0_| = *m*_0_, |*T*_0_| = *n*_0_ and *m*_0_, *n*_0_ = Ω(*n*^ϵ^) *for some ϵ* > 0. *Then for any η* > 1, *as n* → ∞, *we have*

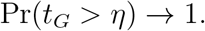

In determining the significance of IGDBs, the simultaneous testing needs to be accounted for all potential IGDBs. Besides, a rejection region (*η*) should be determined based on the distribution of *t*_*G*_ under null model. Hence, we employ the commonly used permutation test procedure in the field of neuroimaging (Zalesky et al., 2010; Nichols, 2012) to empirically approximate the distribution of the likelihood-based statistic *t*_*G*_ under the IGDB null and control the family-wise error rates (FWER). We describe the detailed testing procedure in the Web Appendix A. The p-values of multiple IGDBs can be observed by considering each IGDB individually.

## 5 Results

We applied the IGDB approach to the motivating data set. The FA measures of DTI at 117,139 voxels were used in this study to characterize the white matter integrity (Kochunov et al., 2015, 2016). The image acquisition parameters are described in the Web Appendix B. Regarding genetic variants, 10,595,779 SNPs passed the quality control filters in HCP data set (MAF<0.01; HQE<1e-6; r-squared>0.03; call rate>0.95) after imputation on the Michigan Imputation Server Minimac3 (https://imputationserver.sph.umich.edu) using the 1000 Genomes Project (phase 1 v3) reference set (Das et al., 2016).

We preprocessed the diffusion weighted images following the ENIGMA-DTI workflow (http://enigma.ini.usc.edu/protocols/dti-protocols/). We further applied the Sequential Oligogenic Linkage Analysis Routines (SOLAR)-Eclipse software (https://www.nitrc.org/projects/se_linux) for the heritability analysis, of which imaging voxels were kept with significant heritability, based on the Fast and Powerful Heritability Inference (FPHI) function of SOLAR-Eclipse (p<0.05) in both the HCP and Amish Connectome Project (ACP). For these voxels, we performed vGWAS while adjusting covariates including sex, age, BWI, and population characteristics using the first 10 principal components in our application. We then performed sure independence screening on SNPs with multiple imaging responses through a direct extension of univariate screening procedure (Zou et al., 2021). 13,498 SNPs survive into further analysis. The details are described in the Web Appendix B.

We tested the imaging-genetic associations between SNPs across 22 chromosomes and voxel-level imaging traits using our proposed method. Based on the procedures described in section 4.1 and 4.2, we extracted IGDBs and performed permutation tests to determine its statistical significance while controlling family-wise error rate (*q* < 0.05). We observe different brain areas being influenced by distinct genetic loci. A Manhattan plot for all SNPs across 22 chromosomes with selected imaging-genetic associations highlighted and tables for snp and voxels across all 22 chromosomes are included in the Web Appendix B.

In this section, we focus on SNPs on chromsome 1 to demonstrate their systematic association patterns with voxel-traits, and then annotate the genes in the detected IGDB. Based on the matrix of association strength ***W***_1178×29627_ (i.e., Figure 3 (a)), we detected an IGDB with 384 SNPs and 3803 voxels as Figure 3 (b) by maximizing the objective function (4). The computation is efficient, which took 20 minutes on a PC with an i7 CPU 3.60 GHz and 64GB memory. We further calculated the *p* value for the IGDB statistical inference via the permutation test, which results in a significant existence of an IGDB with *p* value < 0.001. Although the IGDB is an irreducible subgraph, it can be further refined based on data-driven algorithms and spatial information of imaging data. We applied the existing community detection algorithms (Chen et al., 2018) on similarity matrices observed from the detected IGDB. The refined pattern in Figure 3 (c) displays 6 distinct SNP-voxel association clusters. Note that the refined structure can not be identified without revealing the IGDB by the proposed algorithm.

**Figure 1:**
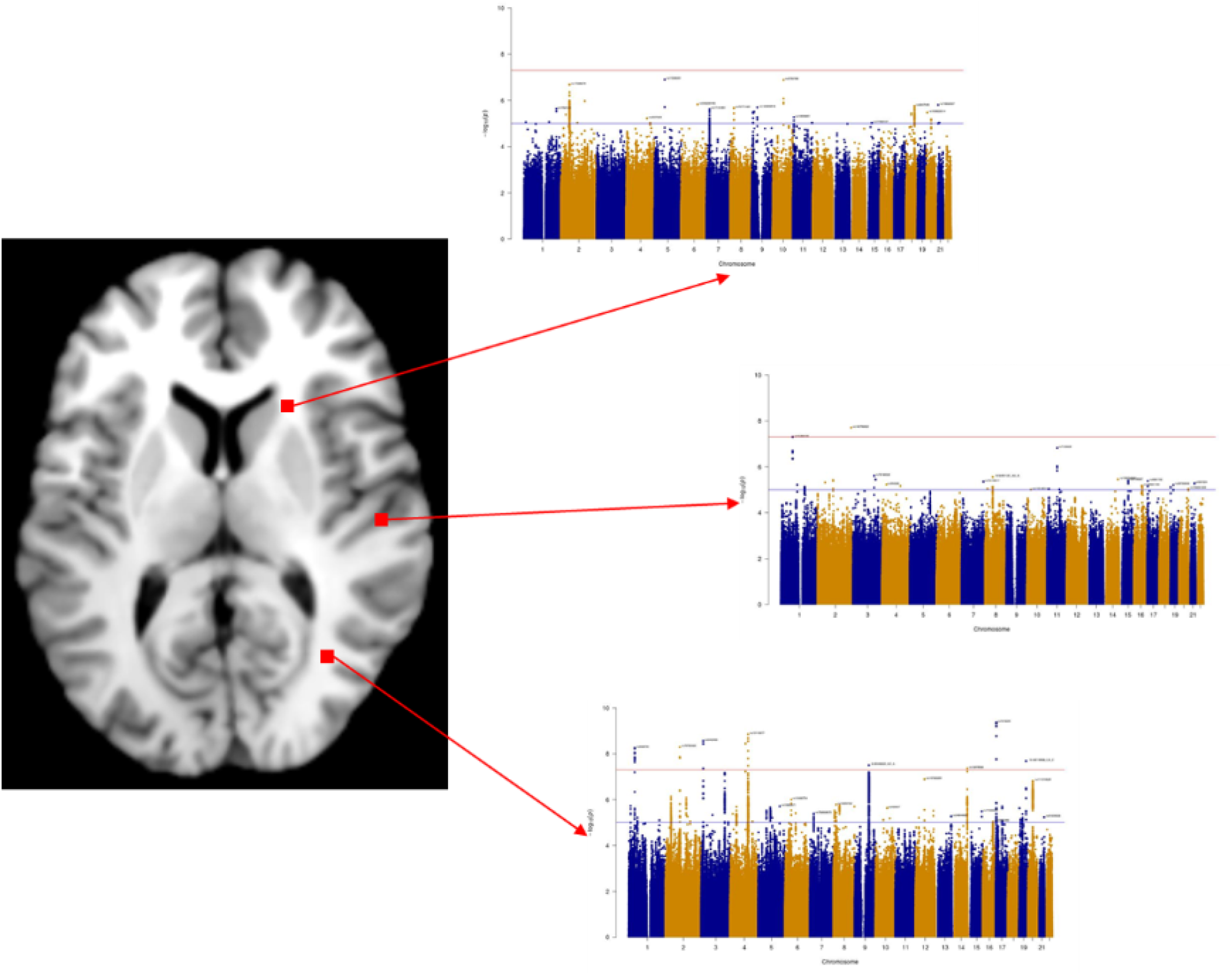
Data structure for vGWAS

**Figure 2:**
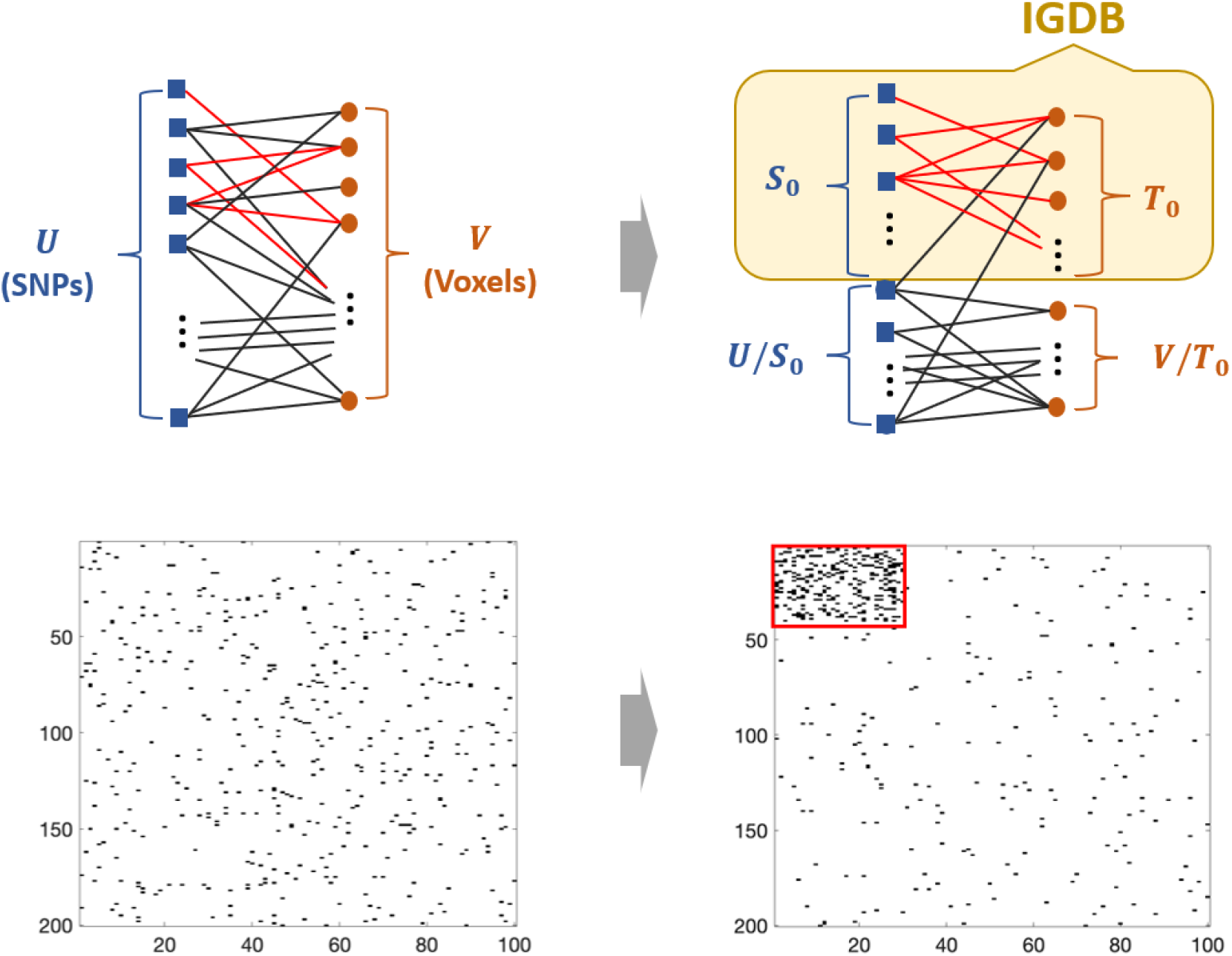
Illustration of a bipartite graph with IGDB structure *G*[*S*_0_, *T*_0_]. The right subfigure highlights *G*[*S*_0_, *T*_0_] in *G* with nodes reordered.

**Figure 3:**
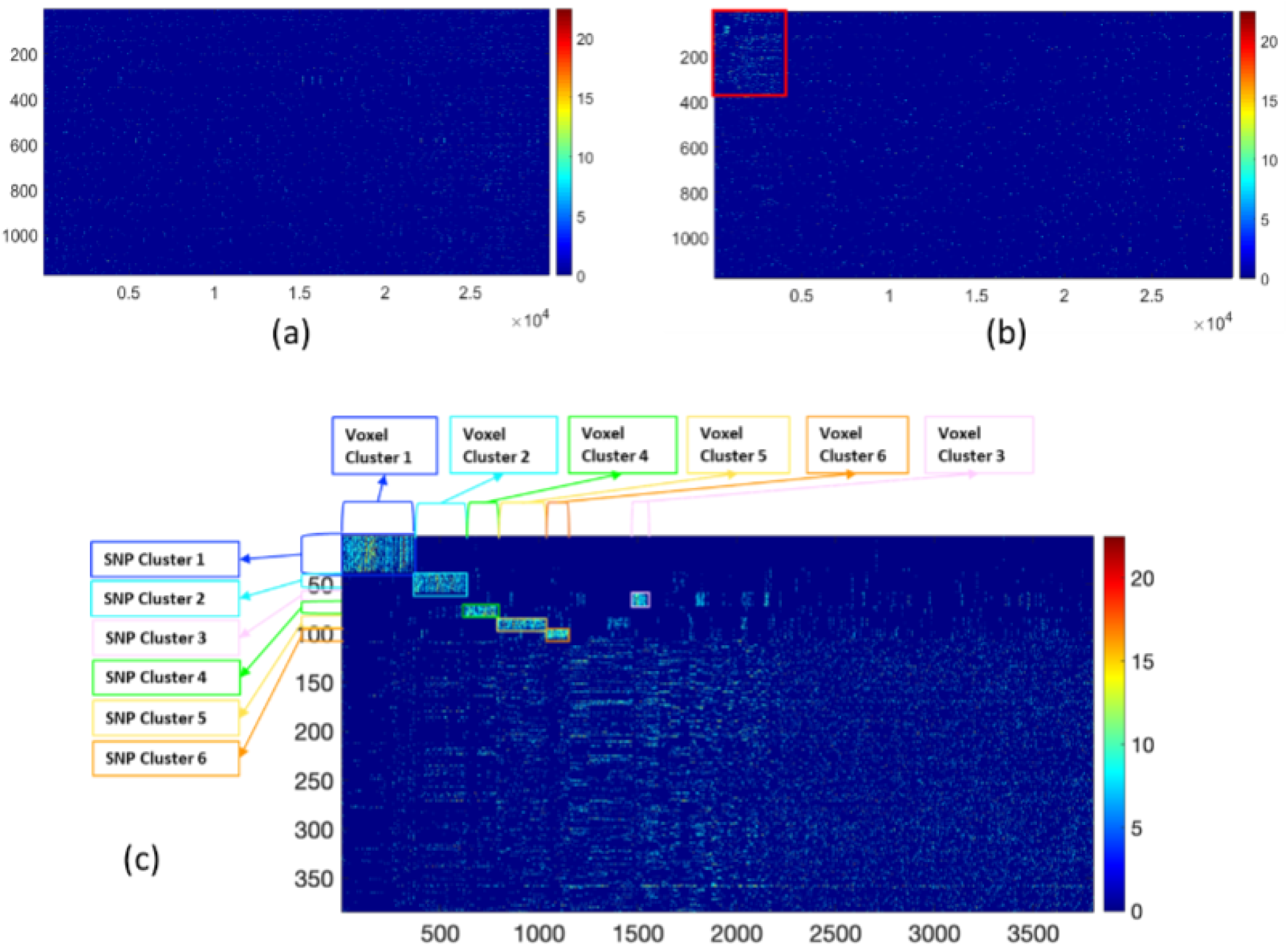
IGDB procedure on chromosome 1: (a) is the input matrix ***W***; (b) demonstrates the detected IGDB; (c)displays the refined pattern of the IGDB

We illustrate the voxel clusters and corresponding SNP sets in Figure 4. For example, the voxel cluster 2 (colored cyan) includes voxels mainly from the splenium of corpus callosum (SCC), part of one of the largest white matter tracts that connects many parts of the brain, and which lesions to often result in many varied neurological issues (Park et al., 2014). To annotate the SNPs in the identified clusters, we queried the SNPs in the QTLbase (http://mulinlab.org/qtlbase/index.html, (Zheng et al., 2020)) for potential expression quantitative trait locus (eQTL) and examined the genes being regulated by these variants in a tissue-specific pattern. The summary of associated genes related with brain tissues is displayed in Web Table 4 as supporting information. In cluster 1, multiple SNPs are linked with the LEPR gene, a protein coding gene for leptin receptor generation that has been shown to be associated with obesity. It has been known the white matter integrity is highly associated with obese disorder and body mass index (Verstynen et al., 2012). Therefore, this cluster reveals the marginal association of (obesity-related) LEPR gene and white matter integrity. In clusters 2, 3, 4, and 5, the associated genes, for example, S100A1, TAF1A, CFH, CFHR3, and DPH5 are associated with immune system functions (http://immunet.princeton.edu/, https://www.innatedb.com/moleculeSearch.do). White matter integrity can be influenced by the immune system functions and systematic inflammation. In cluster 6, the NOS1AP gene has been found to be associated with white matter microstructure in previous studies (Zhao et al., 2019). In addition, the NOS1AP gene is identified to be a risk factor for schizophrenia (Brzustowicz et al., 2004), while the alterations of white matter integrity for patients with schizophrenia were studied in Kubicki et al. (2005). In summary, our findings provided insights into the complex neurogenetic mechanisms of how genetic variants influence imaging traits in a systematic fashion potentially via regulating gene expression and generated hypotheses to be further confirmed in future multi-omics studies.

**Figure 4:**
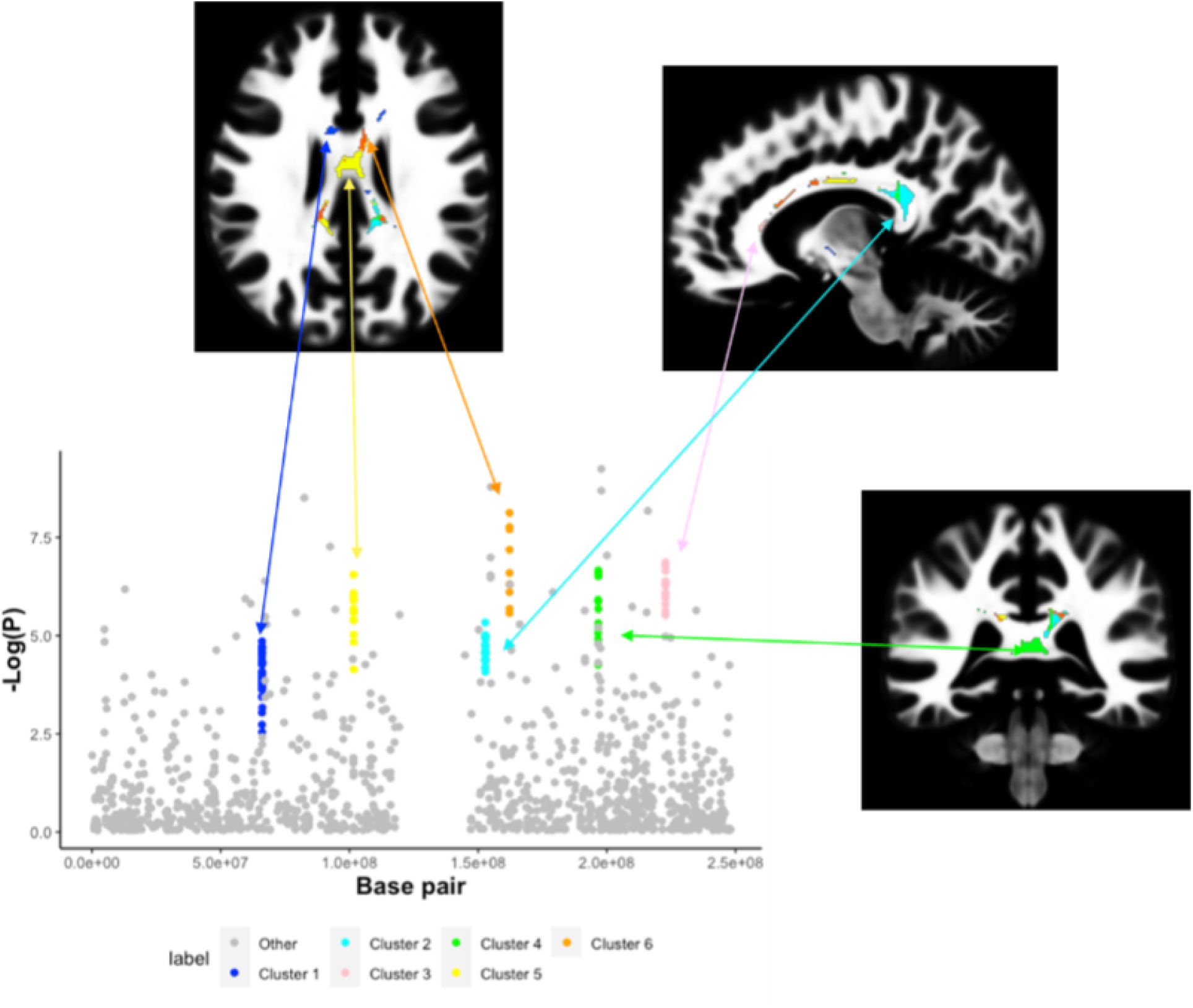
An illustration of the association patterns between SNP and voxel clusters on chromosome 1.

## 6 Simulation Studies

### 6.1 Synthetic data

We evaluate the finite-sample performance of our proposed method based on simulation studies. We generate the input matrix ***W***_*m*×*n*_ based on the two sets of multivariate variables representing genetic variants ***X***_*m*×*L*_ and imaging voxels ***Y***_*n*×*L*_. We let the pattern of ***W***_*m*×*n*_ be determined by a graph *G* = (*U, V, E*). Specifically, we assume there exists an IGDB *G*[*S*_0_, *T*_0_] = (*S*_0_, *T*_0_, *E*[*S*_0_, *T*_0_]) with higher proportion of edges as significant imaging-genetics associations (i.e., *μ*_1_) than the rest of graph (i.e., *μ*_0_). Then, we let the entries of *W*_*m*×*n*_ follow mixture distributions according to *G* as *w*_*uv*_|*δ*_*uv*_ = 1 ~ *μ*_1_*t*_*df*_ (*ν*) + (1 − *μ*_1_)*t*_*df*_ (0), *w*_*uv*_|*δ*_*uv*_ = 0 ~ *μ*_0_*t*_*df*_ (*ν*) + (1 − *μ*_0_)*t*_*df*_ (0), where *δ*_*uv*_ is an indicator variable with *δ*_*uv*_ = 1 for edges in the IGDB and 0 otherwise. *t*_*df*_ (*ν*) and *t*_*df*_ (0) are the non-null and null distributions of imaging-genetics associations respectively. *t*_*df*_ (*ν*) is a *t* distribution with the degree of freedom *L* − *p* (*p* covariates) and non-central parameter 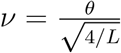 , where *θ* is standardized effect size (e.g., Cohen’s d). *μ*_1_ and *μ*_0_ are the proportions of the non-null distribution within the IGDB and otherwise. We use *m* = 200, *n* = 100, and *L* = 60. We simulate data sets with multiple settings by varying the size of IGDB (i.e., (|*S*_0_|, |*T*_0_|) = (50, 40) and (30, 20)), standard effect size (i.e., *θ* = 0.8, 1, and 1.2), and proportions of noisy edges (i.e., (*μ*_1_, *μ*_0_) = (0.8, 0.2) and (0.9, 0.1)). Additional simulation settings with larger graph and sample sizes are included in the Web Appendix B.

### 6.2 Performance metrics and results

We evaluate the performance of proposed method at two levels. At the subgraph-level, we assess the accuracy of IGDB inference by examining if we can reject the null (i.e., no systematic imaging-genetics association). At the edge-level, we evaluate the accuracy of detected IGDB by comparing it with ground truth in terms of edge differences.

For IGDB inference, we consider a detected IGDB 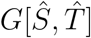 is a recovery of the underlying IGDB *G*[*S*_0_, *T*_0_] if it is rejected in the proposed likelihood-ratio test and has high similarity with *G*[*S*_0_, *T*_0_]. Specifically, we consider 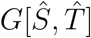 is a true positive detection of *G*[*S*_0_, *T*_0_] if *J*_***X***_ ∧ *J*_***Y***_ is no less than the cutoff with

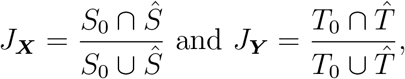

and we succeed to reject the IGDB null hypothesis in the permutation test. We display the results with cutoff 0.8 and 0.9. Therefore, the detected IGDB leads to a false negative finding if the *p*-value in the permutation test is not lower than the a significant level (i.e., 0.05). Besides, we observe a false positive error if 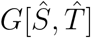 has low similarity to *G*[*S*_0_, *T*_0_] even we rejected the IGDB null hypothesis. We report the accuracy of inference by False Positive Rate (FPR) and False Negative Rate (FNR) among replications.

Furthermore, we compare IGDB to commonly-used multivariate testing methods at the edge-level: positive false discovery rate (pFDR) by Storey (2002) and Bonferroni correction. These correction methods are commonly used in GWAS and vGWAS analysis in practice. We evaluate the true **Δ** = {*δ*_*uv*_}_*u*∈*U,v*∈*V*_ with estimated 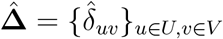 from varied methods. For the proposed method, we obtain the 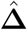 based on the extracted IGDB 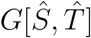 and the hypothesis testing. Particularly, if we reject the IGDB null hypothesis with a detected IGDB 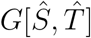, we let 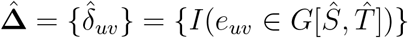. In the case that we fails to reject, we consider 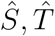 as empty sets such that 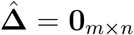. The FDR threshold of 0.2 and corrected *α* level of 0.05 are used in the pFDR and Bonferroni correction respectively.

Subsequently, based on the 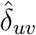 observed from different methods, and true parameters *δ*_*uv*_, we calculate true positive rate (TPR) and true negative rate (TNR) as:

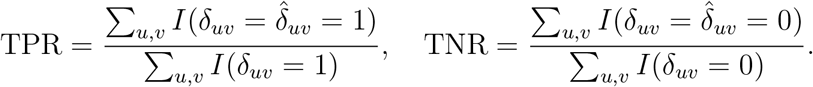

The associated means and standard deviations are reported based on 100 replications for each simulation scenario.

The results from the IGDB inference are summarized in Table 1. The power of the IGDB inference relies on the size and SNR (by different standard effect sizes) of the underlying IGDB *G*[*S*_0_, *T*_0_], which concurs with our theoretical results. We fails to reject the IGDB null hypothesis for one simulated data set with a smaller size (30, 20) and effect size 0.8, and higher noise (0.8, 0.2).

**Table 1:**
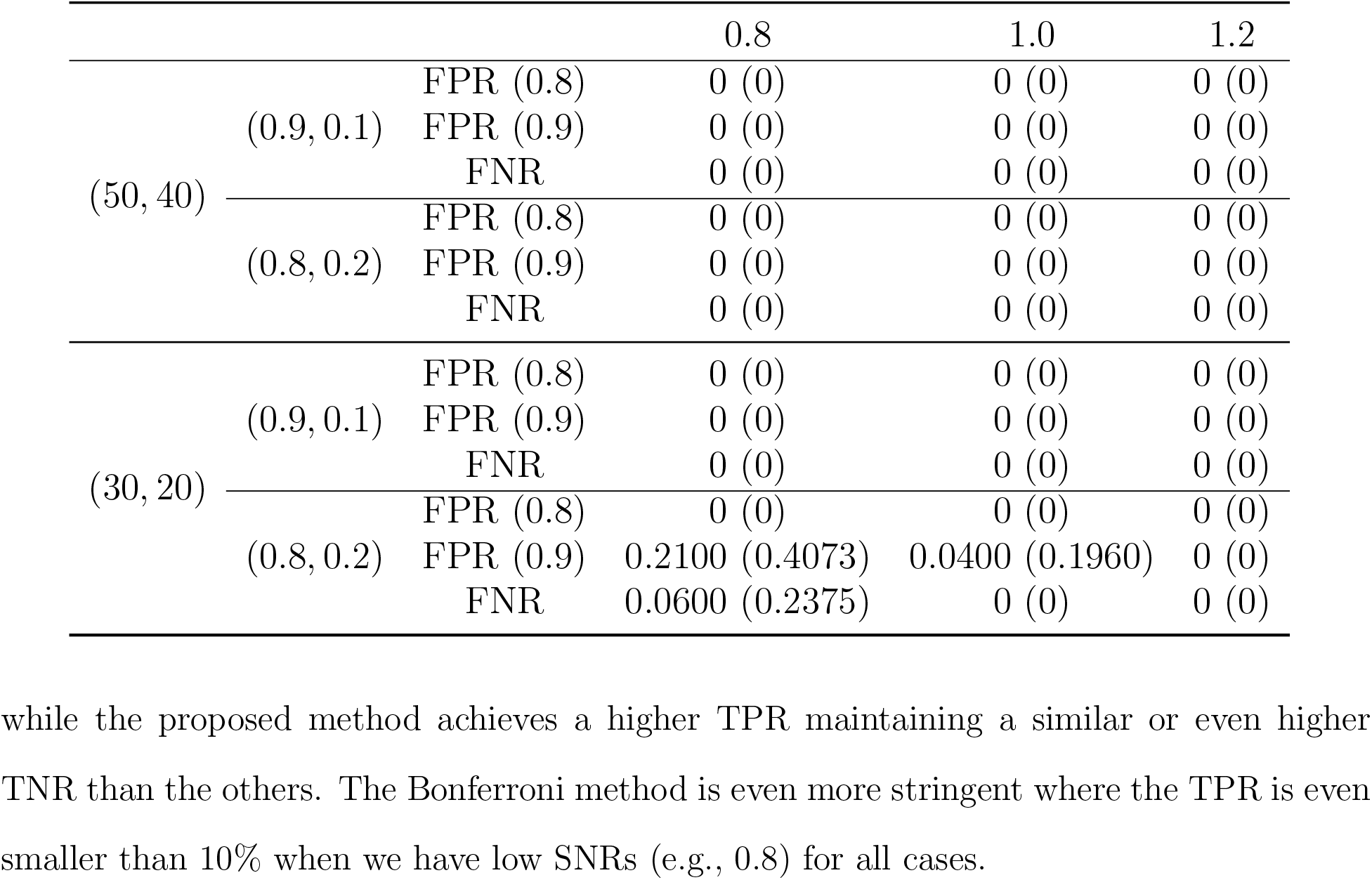
IGDB inference results under varied SNRs and noises

The comparative edge-level results from the proposed method and competing methods are displayed in Table 2 for different sizes of IGDB. All three methods have improved performance with higher SNRs and lower noise levels. The proposed method outperforms pFDR and Bonferroni correction methods for both TPR and TNR under different scenarios. Both pFDR and Bonferroni methods have high TNR but low TPR indicating a stringent cutoff, while the proposed method achieves a higher TPR maintaining a similar or even higher TNR than the others. The Bonferroni method is even more stringent where the TPR is even smaller than 10% when we have low SNRs (e.g., 0.8) for all cases.

**Table 2:**
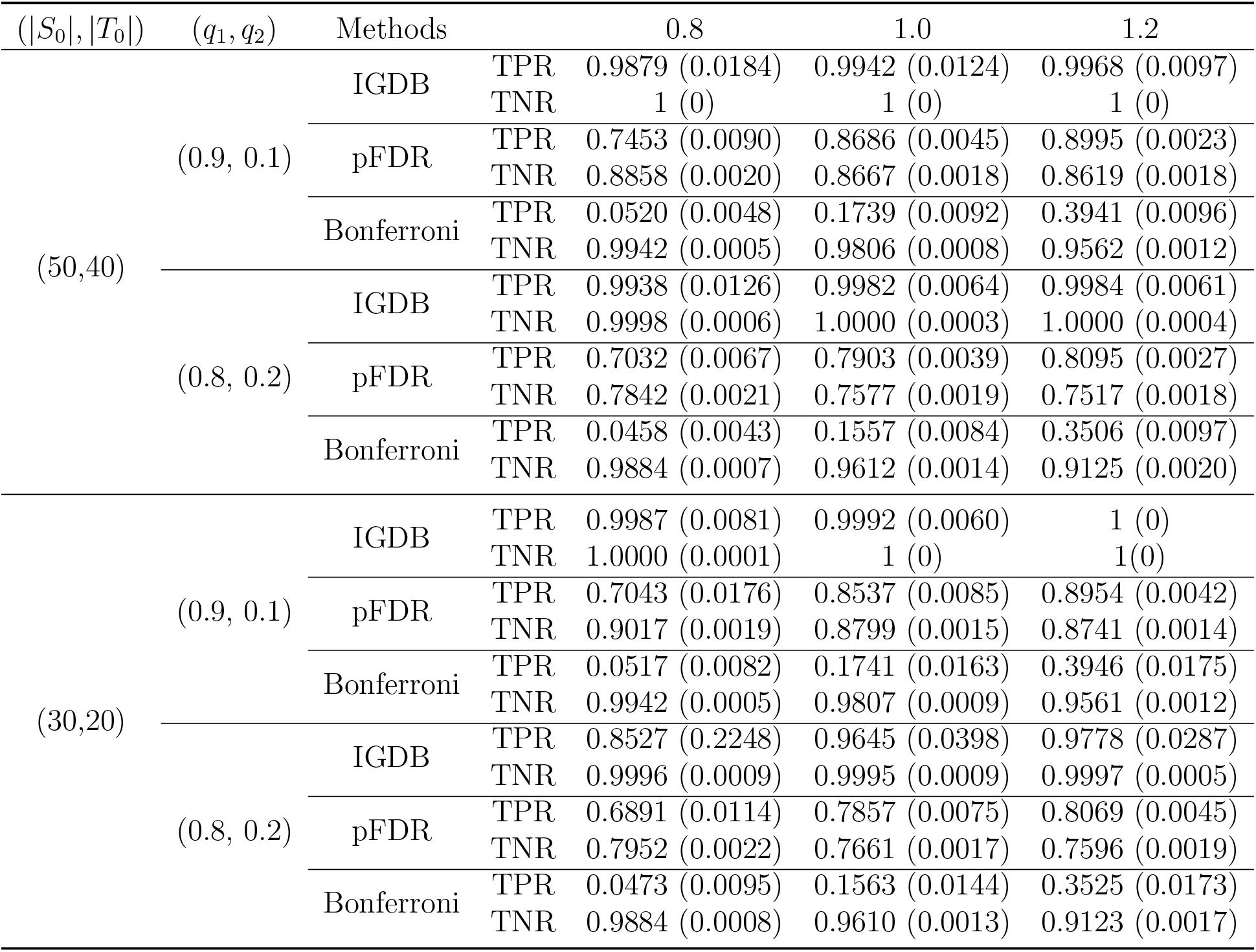
Edge-wise accuracy under varied IGDB sizes, SNRs and noises.

## 7 Discussion

We have developed an IGDB mulivariate to multivariate analysis tool to identify systematic associations between multivariate voxel-level imaging features and multivariate genetic variants. Our method focuses on the systematic polygenic and pleiotropic patterns rather than individual pairwise associations, and thus mitigates the challenges of ultra-high dimensionality due to multivariate to multivariate association analysis.

We develop a new optimization solution to extract IGDB by leveraging its graph properties that we discovered in theoretical study. Our IGDB extraction algorithm is computationally efficient and scalable. The input data for our method could either individual-level or GWAS summary statistics. The IGDB inference method controls the family-wise error rate for IGDB-level findings. We provide theoretical results to guarantee the numerical performance of IGDB extraction and accuracy of the inference model. In real data applications, we identify significant IGDBs where voxels are spatially contiguous and SNPs are functionally correlated confirmed by eQTL. Our IGDB algorithm can also be extended to further constrain the IGDB structure by leveraging the functional annotation of genetic variants (Li et al., 2020).

